# HIV-1 Infection in Humanized Microglial Mice Disrupts Feeding Behavior and Circadian Rhythms with Cortical Neuroinflammation

**DOI:** 10.64898/2026.06.25.733229

**Authors:** Amanda J. Fernandes, Edward Makarov, Saumi Mathews, Debashis Dutta, Matthew Thiele, Mystera M. Samuelson, Santhi Gorantla

**Affiliations:** Department of Pathology and Microbiology, College of Medicine, University of Nebraska Medical Center, Omaha, NE 68198; Department of Pharmacology and Experimental Neuroscience, College of Medicine, University of Nebraska Medical Center, Omaha, NE 68198; Department of Environmental, Agricultural & Occupational Health, College of Public Health, University of Nebraska Medical Center, Omaha, NE 68198

**Keywords:** Humanized mice, NeuroHIV, Home cage behavior, Circadian rhythm, Feeding behavior, Neuroinflammation, Neuropathology

## Abstract

Although effective antiretroviral therapy (ART) has substantially reduced the severity of human immunodeficiency virus (HIV)-associated neurocognitive disorders (HAND), the condition remains highly prevalent. Understanding HAND has been a challenge due to the lack of small animal models capable of supporting productive HIV infection in the brain. Recent advances in humanized mouse models with engrafted human glial cells now enable systemic HIV infection that extends to the central nervous system, offering a powerful platform to study HAND pathogenesis. In this study, we investigated behavioral alterations and neuropathological changes associated with HIV infection. Using home-cage monitoring, we observed that HIV-infected mice exhibited reduced feeding efficiency, consuming less food despite spending more time at the feeder, compared to uninfected controls. Additionally, infected animals displayed disrupted circadian rhythms, with a significant correlation between central nervous system viral load and increased locomotor activity during the light cycle. Neuropathological analyses revealed region-specific vulnerability, with the cortex exhibiting pronounced inflammatory and neurodegenerative changes. These findings were supported by transcriptomic profiling, which demonstrated heightened inflammatory and antiviral gene expression in the cortex, along with differentially expressed genes associated with neuropathology and behavioral deficits. Together, these results highlight distinct region-specific responses to HIV infection in the brain and establish this humanized mouse model as a valuable tool for elucidating the mechanisms underlying HAND and its associated behavioral deficits.

## 1. Introduction

Although currently there is no cure or vaccine for HIV, antiretroviral therapy (ART) does prevent mortality and extend life by disrupting the HIV viral life cycle and suppressing replication of the virus (van Heuvel et al., 2022). Despite the effectiveness of ART, nearly 50% of people living with HIV (PLWH) suffer from a spectrum of neurological complications, ranging from minor neurocognitive dysfunction to severe dementia, which is collectively defined as HAND (McArthur et al., 2010). While ART has reduced the incidence of the most severe forms of this disorder, the milder forms of HAND still show a high frequency of incidence. Therefore, HAND continues to significantly impact PLWH, affecting quality of life, daily functioning, and independence (Heaton et al., 1994; Saylor et al., 2016) and persists even in individuals receiving suppressive ART, likely reflecting ongoing CNS immune activation and viral reservoir activity. Beyond cognitive impairment, HAND is frequently accompanied by a range of neuropsychiatric and behavioral disturbances that substantially compound disease burden (Fatokun et al., 2025; Sukumaran & Sabin, 2023). Individuals with HAND often exhibit deficits in attention, executive function, processing speed, and memory, which translate into difficulties in medication adherence, employment, and social interactions (Hinkin et al., 2004; Kelly et al., 2014; Moschopoulos et al., 2024). In addition, behavioral symptoms such as apathy, irritability, and impaired decision-making are common, further reducing functional independence (Moschopoulos et al., 2024; Singer & Thames, 2016; Thompson et al., 2024). Psychiatric comorbidities, including depression, anxiety, and substance use disorders, are highly prevalent among PLWH (Chander et al., 2006; Gaynes et al., 2008; Hoare et al., 2021). These conditions are associated with poorer clinical outcomes, increased risk of ART non-adherence, and accelerated disease progression, highlighting the complex bidirectional relationship between neurocognitive impairment and mental health (Blashill et al., 2015; Kidie et al., 2026). Importantly, despite the high prevalence and significant clinical impact of HAND, there are currently no approved or widely effective therapies specifically targeting its underlying neuropathogenesis (Bougea et al., 2019). Existing management strategies primarily rely on optimized ART and symptomatic treatment of psychiatric manifestations, which do not adequately address persistent neuroinflammation, viral reservoirs in the central nervous system, or ongoing neuronal injury (Ellis et al., 2014; Gendelman & Gelbard, 2014; Kolson, 2022). This unmet therapeutic need underscores the urgency of developing targeted interventions to mitigate neurocognitive decline, improve neuropsychiatric outcomes, and enhance overall quality of life in PLWH (Scanlan et al., 2022).

The development of effective therapeutics for HAND has been particularly challenging, in large part due to the lack of small, biologically relevant animal models that faithfully recapitulate key aspects of HIV neuropathogenesis (Gorantla, Poluektova, et al., 2012; Tyor & Bimonte-Nelson, 2018). HIV-1 exhibits strict species tropism and does not naturally infect rodent cells, thereby limiting the utility of conventional mouse models for studying viral replication, persistence, and associated neurocognitive impairments (Honeycutt et al., 2015). As a result, early models relied heavily on transgenic expression of viral proteins or peripheral immune activation, which, while informative, fail to fully capture the complexity of productive infection, viral latency, and the dynamic interactions between infected human immune cells and the central nervous system (CNS) (Langford et al., 2018; Thaney et al., 2018). This limitation has hindered mechanistic studies of HAND, particularly those aimed at understanding how HIV establishes and maintains latent reservoirs within the brain, in microglia and infiltrating macrophages (Jaeger & Nath, 2012; Waight et al., 2022). Moreover, the absence of robust small animal models that exhibit both systemic infection and CNS involvement has constrained preclinical testing of candidate therapeutics targeting viral reservoirs, neuroinflammation, and synaptic dysfunction (Omeragic et al., 2020; W. Ru & S. J. Tang, 2017). While non-human primate models, such as simian immunodeficiency virus (SIV)-infected macaques, have provided valuable insights, their high cost, ethical considerations, and limited scalability restrict their widespread use for therapeutic screening (Garcia-Tellez et al., 2016).

Recent advances in humanized mouse models, particularly those engrafted with human immune cells, have enabled important studies of HIV infection and antiretroviral drug efficacy (Honeycutt et al., 2015; Marsden, 2020; Nischang et al., 2012; Victor Garcia, 2016). However, these models largely lack bona fide human microglia in the mouse brain (Evering & Tsuji, 2018; Gorantla, Gendelman, et al., 2012; Honeycutt et al., 2015). Instead, they exhibit only sparse infiltration of human myeloid cells, primarily macrophages localized to the meninges and, to a lesser extent, perivascular spaces, with very limited presence within the brain parenchyma (Gorantla et al., 2010). Consequently, HIV infection within the central nervous system (CNS) is minimal and does not adequately recapitulate the cellular and anatomical context of human brain infection (Boska et al., 2014). To address these limitations, we recently developed a humanized mouse model that supports robust reconstitution of human microglia within the CNS (Mathews et al., 2019). This was achieved through transgenic expression of human interleukin-34 (hIL-34) in NOD/Shi-scid/IL-2Rγnull (NOG) mice, a key cytokine required for microglial development, survival, and maintenance (Mathews et al., 2019; Zhang et al., 2021). In this model, transplantation of human hematopoietic stem cells results in efficient engraftment of human myeloid lineage cells, including microglia-like cells that extensively populate the brain parenchyma. These cells exhibit morphological, phenotypic, and functional characteristics consistent with primary human microglia, thereby providing a more physiologically relevant platform for studying HIV neuropathogenesis. Importantly, this model supports systemic HIV infection following peripheral inoculation, with subsequent viral dissemination to the brain, where productive infection of microglia is established. Infected microglia exhibit active viral replication and induction of human specific interferon and antiviral responses, recapitulating key innate immune features observed in PLWH.

To model HIV-driven neurobehavioral and neuroinflammatory outcomes, we performed longitudinal behavioral assessments by monitoring animals in their home-cage environment, enabling unbiased detection of alterations in activity, exploration, and circadian behavioral patterns. In parallel, we conducted neuropathological analyses, including immunohistochemical characterization of brain tissue, along with transcriptomic profiling of brain regions to define HIV-induced neuroinflammatory and neurodegenerative changes. Together, these approaches establish a translational platform that integrates productive HIV brain infection with measurable behavioral, cellular, and molecular outcomes. Collectively, this model provides a comprehensive system to dissect mechanisms of HAND pathogenesis and offers a much-needed preclinical system for the development and evaluation of targeted therapeutics aimed at mitigating HIV-associated neurocognitive impairment.

## 2. Materials and Methods

### 2.1. NOG-hIL-34 colony

Mice are maintained in microisolators under specific pathogen-free conditions. All animal studies were conducted in accordance with protocols approved by the Institutional Animal Care and Use Committee (IACUC) at the University of Nebraska Medical Center (UNMC). The NOD.Cg-*Prkdc^scid^IL2rg^tm1Sug^*Tg (CMV-IL34)1/Jic (NOG-hIL-34) mouse colony, previously described (Mathews et al., 2019), was maintained as a heterozygous colony and bred by pairing NOG-hIL-34 heterozygous female with NOG male mice obtained from Taconic Biosciences, Germantown, NY. Offsprings were genotyped for human IL-34 transgene using genomic DNA isolated from tail biopsies by proteinase K digestion followed by standard purification according to manufacturer’s protocol for genomic DNA isolation (Thermo Fisher, Waltham, MA). The presence of the human IL-34 transgene was determined by TaqMan quantitative PCR using primers specific for the human IL34 sequence (forward: 5′-AGGAGACCTGGACCTGCTAA-3′; reverse: 5′-TGTAGGCGTTGAGGTTGATG-3′) and a FAM-labeled probe (5′-CTGCTGCTGCTGGAGCTGCTG-3′), designed to selectively amplify the transgenic cassette. Amplification reactions were performed in a 20 µL volume containing TaqMan Universal PCR Master Mix and 10–50 ng of genomic DNA under standard cycling conditions (95°C for 10 min followed by 40 cycles of 95°C for 15 s and 60°C for 1 min). Only mice positive for the hIL-34 transgene were included in subsequent human immune cell engraftment and HI infection studies.

### 2.2. Human cell reconstitution of mice

CD34+ hematopoietic stem cells (hu-HSCs) were isolated from the human umbilical cord blood of healthy full-term newborns after obtaining parental consent and approval from the Institutional Review Board (IRB) of UNMC. Magnetic beads tagged with antibodies to human CD34 (Miltenyi Biotec) were used to isolate CD34+ HSC, and the purity was determined to be >90%. The isolated cells were used fresh or stored in liquid nitrogen for further use. Newborn NOG-hIL-34 mouse pups were injected with HSCs intrahepatically 10⁵ cells per pup, four hours after irradiation at 1 Gy (RS 2000 X-ray Irradiator, Rad Source Technologies, Inc., Suwanee, GA, USA)(Mathews et al., 2019). At 15-20 weeks of age, animals were tested for human immune system engraftment using flow cytometry of peripheral blood with a multichromatic antibody panel for human antigens CD45, CD3, CD4, CD8, CD19, and CD14. Mice demonstrating robust human immune cell reconstitution in peripheral blood were selected for subsequent experiments. As reported by Mathews et al. (Mathews et al., 2019), this model supports the development of human microglia-like cells that populate the brain parenchyma, enabling downstream studies of HIV infection and neuroinflammation.

### 2.3. Flow cytometry

Blood samples were collected either from facial submandibular vein or by direct heart puncture at euthanasia, while splenocytes were collected by crushing the spleen tissue through a 40 µm cell strainer (Mathews et al., 2019). Briefly, cells were reconstituted in a fluorescence-activated cell sorting (FACS) buffer (2% fetal bovine serum (FBS) in phosphate-buffered saline (PBS)) and incubated with a cocktail of human cell marker-specific antibodies – CD45+ fluoresceine isothiocyanate (FITC), CD19+ phycoerythrin-cyanin 5 (PE-Cy5), CD14+ phycoerythrin (PE), CD3+ alexa fluor 700 (AF700), CD4+ allophycocyanin (APC), and CD8+ brilliant violet 421 (BV421), for 30 minutes at 4 °C. FACS lysing solution and FACS buffer were used to lyse red blood cells (RBCs) and wash the cells, respectively. All reagents and calibration beads were obtained from BD Biosciences, San Jose, CA. Cells were fixed with 2% paraformaldehyde (PFA), and data were acquired with FACS Diva v6 in a BD LSR2 flow cytometer (BD Biosciences), with data analysis done using FLOWJO analysis software version 10.10.0.

### 2.4. HIV infection

Mice ∼20 weeks of age were intraperitoneally infected with 10^4^ tissue culture infection dose_50_ (TCID_50_) of macrophage-trophic HIV-1_ADA_ strain. Viral stocks were prepared as described previously on human monocyte-derived macrophages (Gendelman et al., 1988).

### 2.5. Isolation of RNA and HIV measurement

Mouse blood was collected post-infection from the facial submandibular vein and by cardiocentesis at euthanasia. RNA from plasma was isolated using the QIAmp® Viral Mini Kit, while RNA from tissues such as the brain and spleen was obtained by isolating RNA using the AllPrep® DNA/RNA Mini Kit (Qiagen, Hilden, Germany). HIV RNA copy numbers were quantified with a digital droplet PCR (ddPCR) assay using a HIV-1 gag TaqMan probe and primers set with the sequences: 5’-FAM-CATCAATGAGGAAGCTGCAGAATGGGATAGA-TAMRA-3’ (Applied Biosystems, Foster City, CA), forward (5’-ACATCAAGCAGCCATGCAAAT-3’) and reverse (5’-ATCTGGCCTGGTGCAATAGG-3’) primers’ nucleotide sequences (Invitrogen, Thermo Fisher Scientific). A QX200 ddPCR system (Bio-Rad, Hercules, CA) was used to determine the HIV RNA copies in plasma and tissue samples. All reagents used for ddPCR were obtained from Bio-Rad. PCR reaction mix containing 200 ng of RNA sample, forward, reverse primers, probe, nuclease-free water, and one-step reverse transcriptase ddPCR (RT-ddPCR) advanced super mix (SM) was loaded into a 96-well PCR plate. An automated droplet generator was used to generate droplets. PCR was performed for reaction samples containing more than 20,000 droplets in 4 steps, with the first step of reverse transcription to synthesize cDNA from the RNA template at 48 °C for 1 hour, followed by initial denaturation at 95 °C for 10 minutes. After this, PCR cycling was with denaturation at 95 °C for 30 seconds, followed by annealing/extension at 60 °C for 1 minute, repeated for 40 cycles. The final extension step was performed at 98 °C for 10 minutes. Following the PCR thermocycling reaction, droplets were read for fluorescence signal positivity in the QX200 droplet reader. HIV gene copies per total assay volume (20 µL) were quantified from raw data by Quantasoft Pro software. Human CD45 levels in tissue RNA were measured using ddPCR with a primer and probe set (assay ID Hs00365634_g1; catalog #4331182; Thermo Fisher Scientific).

### 2.6. Home cage monitoring (HCM)

Details describing the home cage behavioral monitoring system and quality control during data acquisition and analysis have been previously published (Goulding et al., 2008; Knibbe-Hollinger et al., 2015; Parkison et al., 2012). Briefly, mice were housed individually in instrumented cages (48.3 cm x 25.7 cm x 15.2 cm, PC10196HT, Allentown) with ad-lib access to food (while breaking a photobeam) and water (while activating a capacitive lickometer). Mouse position was measured by load cells (LSB200, FUTEK) located in different regions of the cage. Data was acquired in binary form using LabView software (National Instruments, TX) and processed using custom-written MATLAB (MathWorks, MA) code. Mice were fed powdered chow (TekLad, Inotiv, irradiated 7012, Madison, WI) and autoclaved water (prepared in-house). The room maintained a controlled 12-hour light/12-hour dark cycle (lights on from 0600 to 1800 h). Cohorts of uninfected control (n = 11) and HIV-infected (n = 14) mice were evaluated. HIV-infected mice were transferred to the home cage monitoring system at 6 weeks post-infection and continuously monitored for 21 days, including a 5-day habituation period.

### 2.7. Data Validation for Home Cage Monitoring

Validation studies were performed to verify that the number of licks and feeder photobeam break time were reliable indicators of fluid and food consumption, respectively. The last 16 days of data were used for analysis, with the first 5 days for habituation. Due to the large volumes of behavioral data generated, methods to maximize data quality, such as algorithms developed in-house, were used to collect data from each device to cross-check the performance of other devices in each cage and to summarize the data (Goulding et al., 2008).

### 2.8. Immunohistochemical Analyses

Five-µm thick PFA-fixed paraffin-embedded spleen and brain tissues were stained with primary antibodies HLA-DQ/DR/DP (CR3/43) (1:100, Novus Biologicals, Littleton, CO) and HIV-1 p24 (05-001) (1:50, Santacruz Biotechnology, Dallas, TX) after antigen retrieval with Trilogy solution (Sigma-Aldrich, St. Louis, MO). Gene Tex One Step Polymer HRP anti-mouse/rat/rabbit (Irvine, CA) was used for primary antibody detection, and the signal was developed by using SIGMAFAST^TM^ 3, 3’-Diaminobenzidine (DAB) Detection IHC Kit (Sigma Aldrich, St. Louis, MO) as per manufacturer’s instructions. The DAB signal developed was brown, and nuclei were counterstained with Mayer’s hematoxylin.

For immunofluorescence staining, sagittal brain sections were treated with different combinations of primary antibodies, including mouse (Ms) anti-human HLA-DQ/DR/DP (CR3/43) (1:100, Novus Biologicals, Littleton, CO), rabbit (Rb) anti-glial fibrillary acidic protein (GFAP) (1:1000, Dako, Carpentaria, CA), Rb anti-microtubule associated protein 2 (MAP2) (1:500, Millipore, Burlington, MA), Rb anti-synaptophysin (SYN) (YE269) (1:800, Abcam, Cambridge, MA), Rb anti-neurofilament heavy chain (NFL-H) (1:400, Millipore, Burlington, MA), Ms monoclonal myelin oligodendrocyte glycoprotein (MOG) (1:500, Santacruz Biotech, TX), and Rb polyclonal myelin associated glycoprotein (MAG) (1:500, Protein-tech, Rosemont, IL). For secondary antibodies, nuclei were labeled with Alexa fluor 594-conjugated goat anti-Rb IgG (1:200, Invitrogen, Grand Island, NY) and Alexa fluor 488-conjugated goat anti-Ms IgG (1:200, Invitrogen, Grand Island, NY). Slides were cover slipped with Prolong Gold anti-fade reagent with 4, 6’-diaminido-2-phenylindole (DAPI) mounting medium (Invitrogen, Carlsbad, CA) to label nuclei. Bright-field and fluorescence images were captured with a Nuance EX camera fixed to a Nikon Eclipse E800 microscope using Nuance software (Cambridge Research & Instrumentation (CRI), Woburn, MA) with a 20x objective on a Nuance Multispectral Tissue Imaging System (CRI, Woburn, MA).

### 2.9. Image Analyses

Bright-field and fluorescence images were captured with a Nuance EX camera attached to the Nikon Eclipse E800 microscope with a 20x objective using the Nuance multispectral tissue imaging system (Cambridge Research & Instrumentation (CRI), Woburn, MA). Fluorescent images were captured at different brain regions – hippocampus and cerebral cortex for the neuronal and glial markers. For MAG and MOG, regions with high myelin content – cerebellum and corpus callosum were considered. A minimum of four images were obtained for all markers per animal. Staining for all slides in the control (n = 4) and HIV-infected groups (n = 7) was performed together, and the results were averaged from each individual area for each antigen to represent the fluorescence intensity as a single result per mouse for statistical analysis. Data was expressed as average ± standard error mean (SEM) of signal counts relative to fluorescence units / µm^2^ (Boska et al., 2014).

### 2.10. Next Generation Sequencing

Total RNA was isolated from the brain tissue samples using the AllPrep® DNA/RNA Mini Kit mentioned previously. RNA quality was assessed using the RNA integrity number (RIN), and samples with RIN > 7 were processed for library preparation. cDNA library was constructed using TruSeq RNA Sample Prep Kit and sequenced to a depth of 100 base pairs/read, with ≤40 million reads per sample, and then sequenced with HiSeq 2000 system (Illumina Inc., San Diego, CA) (Ping et al., 2014). The sequencing data was filtered with SOAPnuke by (1) removing reads containing sequencing adapter; (2) removing reads whose low-quality base ratio (base quality less than or equal to 15) is more than 20%; (3) removing reads whose unknown base (‘N’ base) ratio is more than 5% (Li et al., 2008). Clean reads were stored in FASTQ format and mapped to the human (GRCh37.p13) and mouse (GRCm38.p6) reference genomes separately using HISAT2 (Kim et al., 2015). For gene identification, Bowtie was applied to align the clean reads to the gene set (Langmead & Salzberg, 2012). Expression level of each gene was calculated by RSEM (v1.2.28) to get read count, FPKM (fragments per kilobase of exon per million mapped reads), and TPM (transcripts per million) (Li & Dewey, 2011). Differentially expressed genes (DEGs) were determined using the R-package DEseq with the criteria of a log2 fold change greater than 1 or less than -1 and p-value < 0.05 (Wang et al., 2010). The complete dataset containing DESeq fold changes and p-values for significant genes is permanently hosted on Zenodo (https://doi.org/10.5281/zenodo.20723232). The upregulated and downregulated genes were selected for pathway and disease analysis using the Ingenuity Pathway Analysis (IPA) (Qiagen Inc., https://digitalinsights.qiagen.com/IPA) (Krämer et al., 2014). Network visualization and analysis were performed using Cytoscape web version 1.0.5 (Shannon et al., 2003).

### 2.11. Statistical analysis

Data were analyzed and plotted using GraphPad Prism 10.0.2 (GraphPad, USA) and expressed as mean ± SEM. Unpaired t-test was performed to compare the two groups using GraphPad Prism software. The *p-*value < 0.05 was considered to indicate a statistically significant difference.

## 3. Results

### 3.1. HIV infection in humanized microglial mice

Humanized microglial mice were divided into two groups: the first group consisted of HIV-infected animals, and the second served as a control group without HIV infection. At ∼20 weeks of age, mice were infected with HIV-1 _ADA_. Blood samples were collected from all animals at regular intervals throughout the study period for HIV RNA measurement and flow cytometry for human immune cells. After confirming the establishment of HIV infection by measuring HIV RNA in the blood, mice were monitored for 21 consecutive days using the home-cage monitoring system (Bonasera et al., 2017; Knibbe-Hollinger et al., 2015). At the end of the study, mice were euthanized, and blood and tissues were collected for viral, immune, histopathological, and transcriptomic analyses. The experimental scheme is illustrated in Fig. 1A. The overall experimental design enabled us to perform an integrative analysis linking the viral profile to the behavioral, histopathological, and transcriptional profiles of these animals.

**Figure 1.**
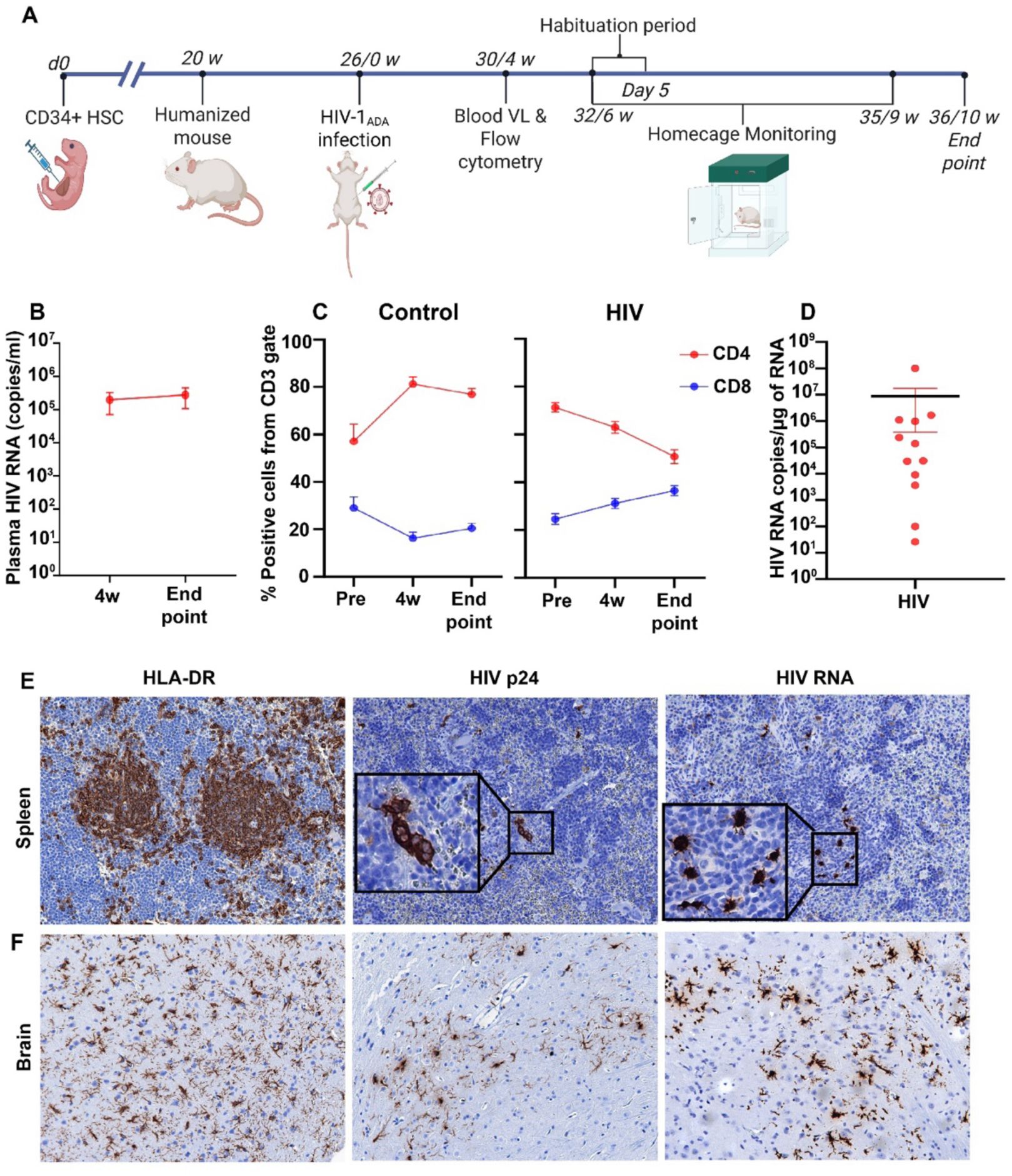
Experimental plan and HIV infection in humanized mice. **(A)** Experimental timeline of immune profile, peripheral and brain viral load, histologic and transcriptomic status of humanized microglial mice (made in Biorender). **(B)** Viral dynamics in plasma of replicate peripheral blood samples from HIV-infected mice at 4 weeks post-infection and end point. **(C)** Flow cytometric analysis of CD4+ and CD8+ T cells in the blood of uninfected and HIV-infected mice before infection (pre), 4-week post-infection, and end point. **(D)** HIV-gag RNA levels in the brain expressed as copies per μg RNA. **(E) and (F)** Representative images of histological analysis of HLADR (for human cells), HIV p24, HIV RNA in the spleens and brains, respectively, from HIV-infected mice.

A progressive decrease in total CD4+ T cells was observed during HIV infection (Fig. 1C). Uninfected controls demonstrated stable CD45 and CD4/CD8 ratios throughout the study (Table S1). All infected animals had sustained viral load (averaged at 2.78 x 10^5^ HIV RNA copies/ml plasma) at the endpoint (Figure 1B). Total HIV-1 RNA in spleen and brain tissues of the infected humanized microglial mice was readily quantified by highly sensitive ddPCR. The brain and spleen had an average of 8.39 x 10^6^ and 3.44 x 10^5^ viral RNA copies per µg of RNA, respectively (Fig. 1D and Fig. S2B). Histological analysis revealed that human leukocytes expressed HIV-1 p24 in the follicular and parafollicular regions of the spleen, as well as human microglia-like cells expressed HIV-1p24 in various brain regions. RNAScope® assay for detecting HIV-1 RNA revealed single brown dots or clusters of dots abundantly at concurrent areas of HIV-1p24 expression (Fig. 1E, F).

### 3.2. HIV infection alters activity patterns, feeding efficiency, and circadian behavior

To characterize the behavioral phenotype of the humanized microglial mice and the impact of HIV infection, home-cage monitoring was performed. Mice were individually housed in the home-cage monitoring system at 6 weeks post-infection and observed for 21 days, including a 5 day habituation period. Parameters measured included resting, movement in place, locomotion, feeding, and drinking behaviors. These behaviors were evaluated based on the light-dark cycle, circadian time, and active and inactive states. Active and inactive states were identified based on position probability density estimates (Goulding et al., 2008; Knibbe-Hollinger et al., 2015; Parkison et al., 2012). Overall, the uninfected control and HIV-infected groups exhibited minimal differences in total locomotion and food/water intake, regardless of the light and dark cycle (Fig. 2A, B, C).

**Figure 2.**
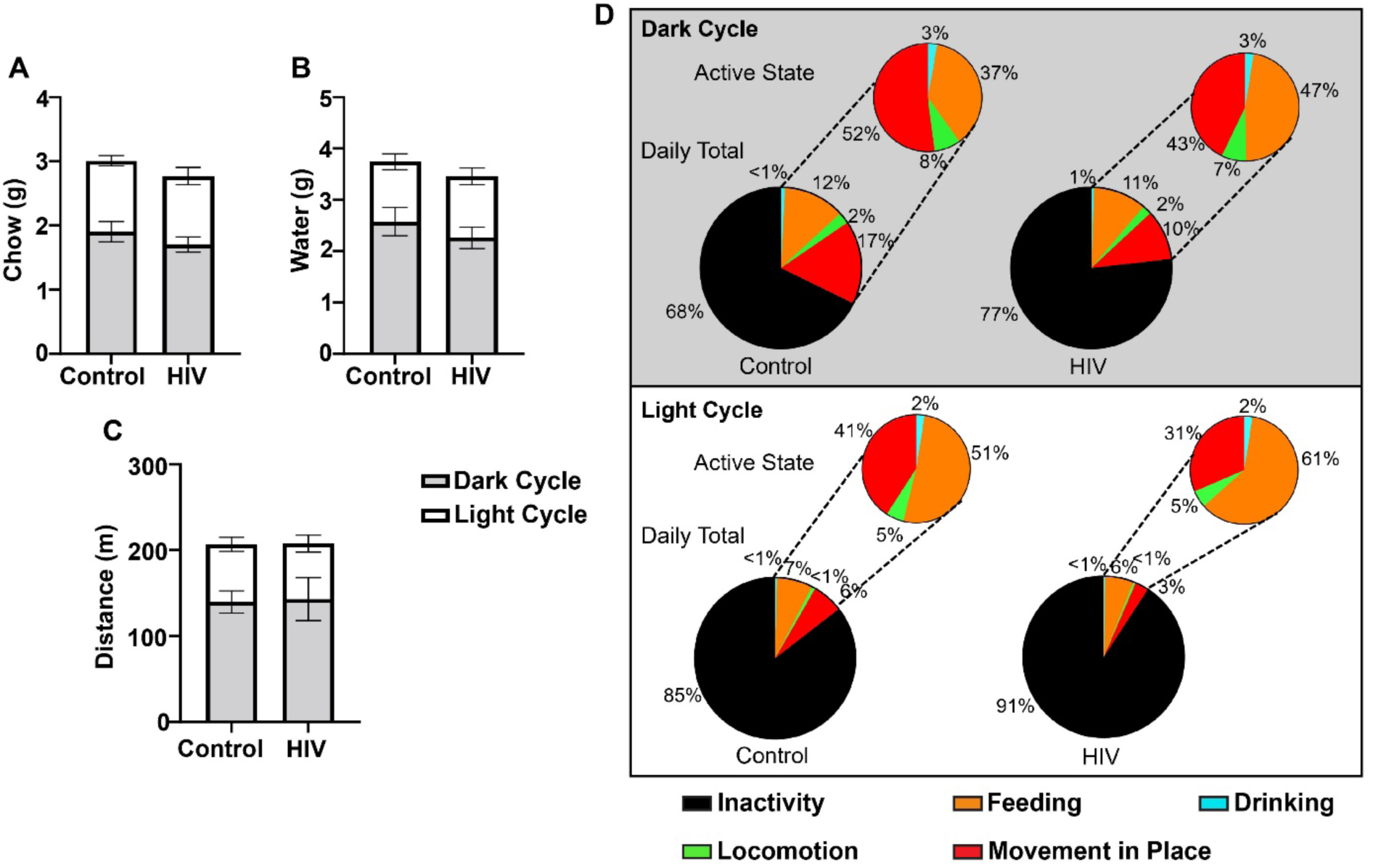
Home cage behavioral analysis. **(A)** Average food intake, **(B)** water intake, and **(C)** overall locomotion for dark cycle (grayed) and light cycle for uninfected controls and infected mice. **(D)** Time budgets analysis comparing daily total and active state time budgets (% of time spent) for uninfected controls and HIV infected mice in the light cycle and dark cycle (grayed). Average food intake, water intake, and overall locomotion are represented as mean ± SEM.

Next, time-budgets analyses were performed to determine the percentage of time spent by these mice in different behaviors both in total day and within the active states during both the light and dark cycles. The active state for mice is defined by behaviors such as foraging and patrolling performed within a regularly traversed home range (Akay et al., 2013; Goulding et al., 2008). Consistent with their nocturnal nature, mice exhibited reduced activity during the light cycle compared to the dark cycle (Fig. 2D). However, during the dark cycle, uninfected control mice displayed a higher frequency of movement-in-place behaviors compared to HIV-infected animals, suggesting increased grooming and/or nest-building activity (Fig. 2D). Importantly, HIV-infected mice spent a greater proportion of time in the inactive state across both light and dark cycles, indicating a possible fatigue-like phenotype associated with HIV-infection. When in the active state, HIV-infected mice were also more likely to engage in feeding behavior than in other activities during both light and dark cycles (Fig. 2D).

When evaluating the relationship between feeding and drinking behaviors, HIV-infected mice exhibited fewer coordinated feeding and drinking bouts, despite spending more time, on average, at the feeder and waterspout than uninfected controls. Compared to HIV-infected mice, uninfected mice were more likely to engage in feeding and drinking bouts in direct succession (Fig. 3A). Average food and water consumption, as well as locomotion, were measured for both control and infected animals in the context of circadian time. Overall, HIV-infected animals displayed a decrease in food and water consumption compared to controls despite a similar peak at the beginning of the dark cycle (Fig. 3B, C). This pattern suggests that, despite spending more time in these behaviors, HIV-infected animals were less efficient at meeting their nutritional and hydration needs than uninfected controls, which achieved similar outcomes with shorter and more frequent feeding and drinking bouts. Control animals exhibited two strong peaks in feeding and drinking behaviors, both at the beginning and end of the dark cycle. In contrast, HIV-infected animals peaked only at the start of the dark cycle (Fig. 3B, C). To assess activity across circadian time, we measured total distance traveled throughout the day. Infected mice did not differ in the extent to which they engaged in active behaviors at the beginning of the dark cycle, in comparison to the control group. However, they exhibited possible fatigue at the end of the dark cycle with a slight decline in activity when compared to controls (Fig. 3D).

**Figure 3.**
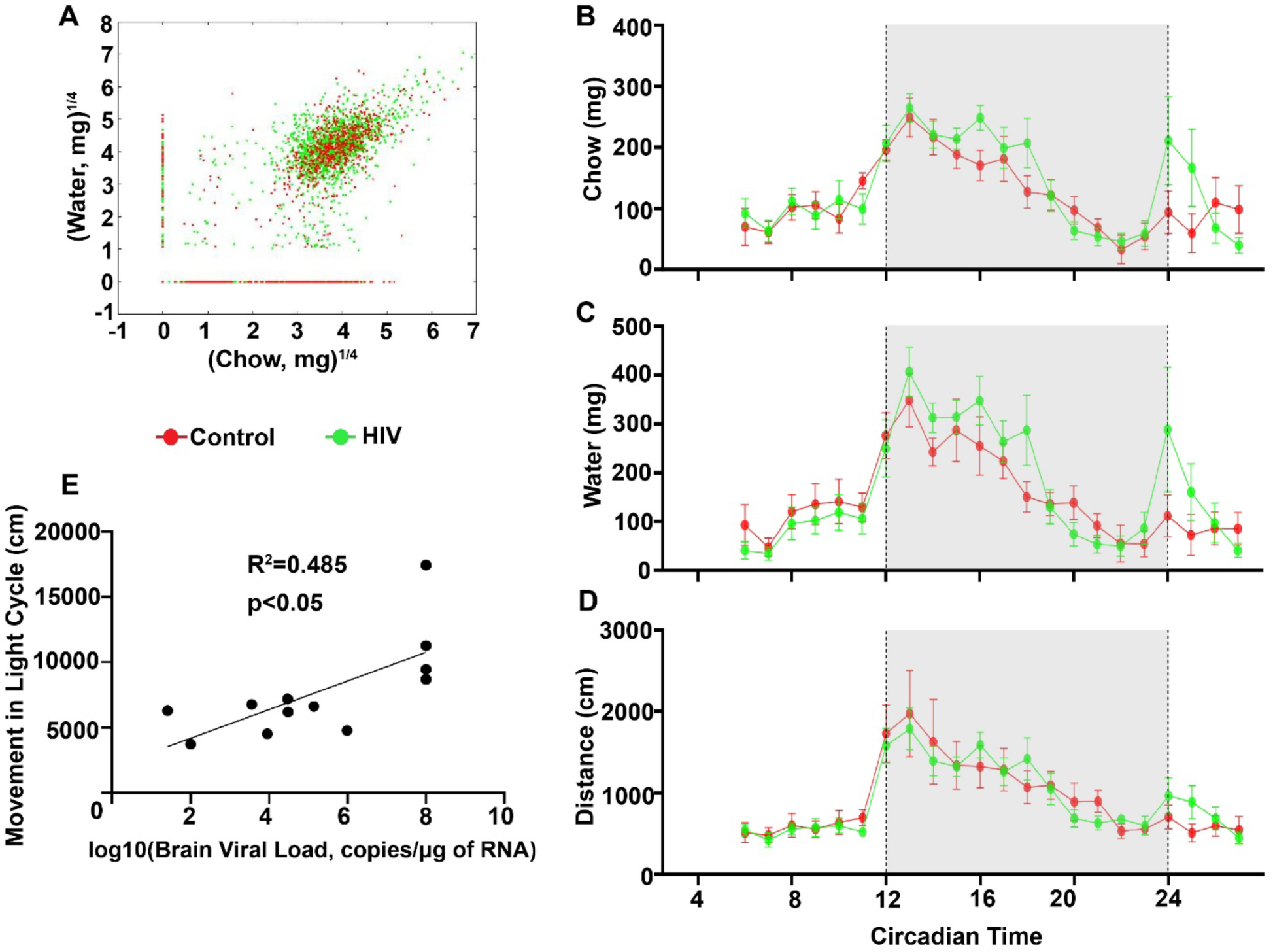
Longitudinal circadian behavioral patterns with HIV-infection. **(A)** Relationship between water and chow consumed by uninfected controls and infected mice. Each dot represents an individual bout. **(B)** Average cumulative food consumption, **(C)** water consumption, and **(D)** locomotion, with respect to circadian time over 16 days. Data is represented as mean ± SEM. The dashed line represents the cut-off from the light-to-dark cycle (grey-shaded area). **(E)** Correlation analysis between brain viral load and movement in the light cycle for infected mice. Controls and infected mice are represented in green and red, respectively.

Correlation analysis of different behavioral parameters and brain viral load showed a statistically significant positive correlation (R^2^=0.485, p<0.05) between brain viral load and locomotion in the light cycle. Due to their nocturnal nature, mice are expected to move less in the light cycle than in the dark cycle. However, in HIV-infected mice, this trend seems to have been reversed and is directly correlated with the increased HIV viral load in the brain (Fig. 3E). Therefore, the home cage monitoring analysis shows that while the uninfected humanized microglial mice behaved similarly to other strains of mice, the HIV-infected group had changes in many behaviors such as activity, appetite, and circadian rhythm due to HIV-infection.

### 3.3. Progressive HIV infection induces neuropathology

To determine whether HIV brain infection in the humanized microglial mice leads to neuropathological changes, we next examined neuronal and glial protein expression across multiple brain regions of uninfected and HIV-infected mice. At the endpoint, brain tissues were collected and processed for histopathological analyses. Sagittal sections of brains were stained for various neuronal and glial markers and analyzed by multispectral fluorescence microscopy. A minimum of two brain sections from each mouse were analyzed, with four representative images per brain region quantified per section. Data were represented as mean fluorescent intensity per μm^2^.

Astrogliosis (activation of astrocytes) was assessed by quantifying GFAP expression. As these cells are the primary glial cell type in the brain, it is essential to measure GFAP expression as a marker of CNS distress and neuroinflammation. When measuring GFAP levels in different brain regions of both uninfected and HIV-infected animals, we observed significantly increased expression in the DG and cortex (p<0.05), indicating neuroinflammation, while we observed an increasing trend in the CA1 region (Figure 4A, B).

**Figure 4.**
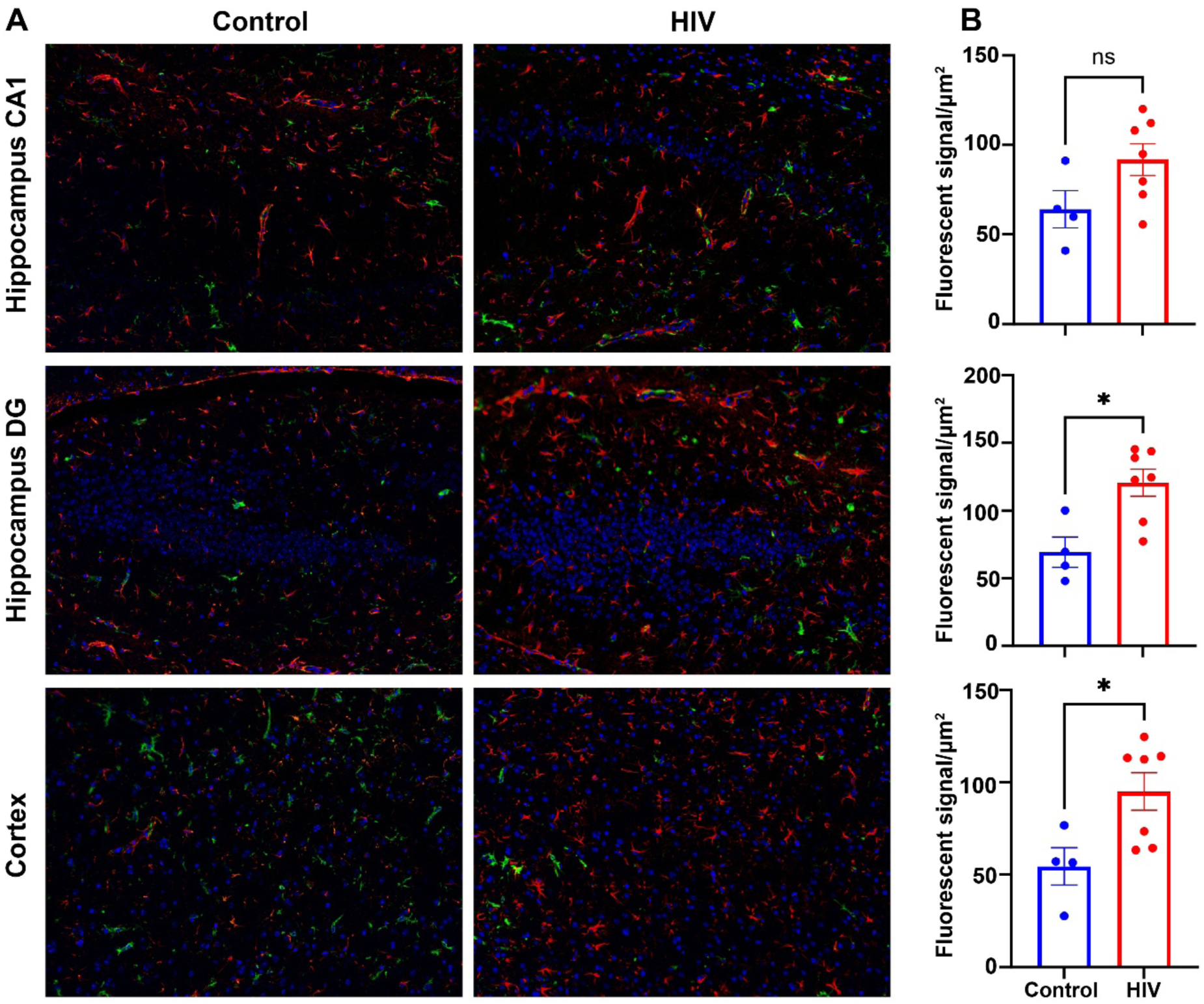
Glial activation in HIV-infected mice. Paraffin-embedded 5 µM brain sections were immunostained with astrocyte marker, GFAP (red), HLA-DR (green), and nuclear counterstain DAPI (blue). **(A)** Representative images captured at 200x original magnification from CA and dentate gyrus (DG) regions of hippocampus, and medial frontal cortex, are shown in uninfected and HIV-infected mice. **(B)** Quantification of fluorescent intensity of GFAP expression in different brain regions was performed using multispectral imaging, comparing HIV-infected and uninfected groups. Data are presented as mean ± SEM, * denotes p<0.05, and ns denotes not significant differences.

To evaluate neuronal injury, we measured synaptic, dendritic, and axonal integrity using presynaptic marker SYN, postsynaptic marker MAP2, and NFL-H protein, respectively. NFL-H expression was significantly reduced across multiple brain regions, indicating neuro-axonal injury, with moderate but significant reductions observed in the hippocampal CA1 and DG (p<0.05) (Figure 5A, D). Synaptic integrity, assessed by SYN immunoreactivity, reveals a punctate and diffuse staining pattern in uninfected brains, whereas HIV-infected mice exhibited a marked reduction in SYN fluorescence intensity. A significant reduction was observed in the medial prefrontal cortex (p<0.001), consistent with presynaptic injury and synaptic dysfunction (Figure 5B, E). Analysis of MAP2 expression revealed a widespread reduction in dendritic integrity across all brain regions examined. The most pronounced decrease was observed in the hippocampal DG (p<0.01), suggesting broader post-synaptic dendritic compromise than presynaptic loss (Figure 5C, F).

**Figure 5.**
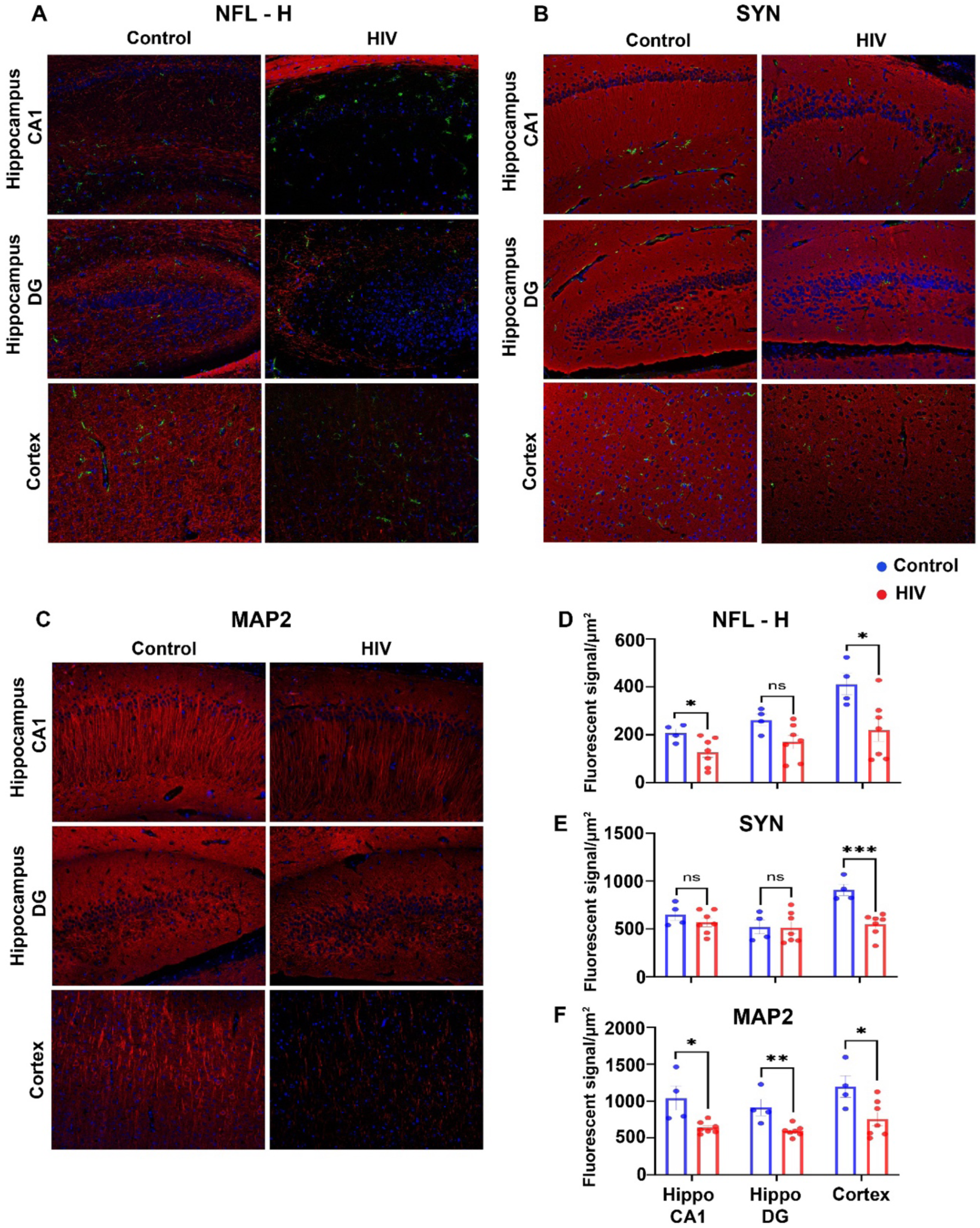
Neuronal injury in HIV-infected mice. Paraffin-embedded (5 µm) brain sections were immunostained for neuronal markers **(A)** neurofilament heavy chain (NFL-H) (red), **(B)** synaptophysin (SYN) (red), and **(C)** microtubule-associated protein-2 (MAP2) (red), together with the microglial activation marker HLA-DR (green), and nuclei were counterstained with DAPI (blue). **(A–C)** Representative images acquired at 200x original magnification from the hippocampal CA and DG, and medial frontal cortex are shown for uninfected and HIV-infected mice. **(D–F)** Quantification of fluorescence intensity for NFL-H, SYN, and MAP2 across different brain regions was performed using multispectral imaging, comparing HIV-infected and uninfected groups. Data are presented as mean ± SEM; *p < 0.05, **p < 0.01, ***p < 0.001, and ns indicates non-significant differences. Quantification of fluorescence intensity across brain regions was performed using multispectral imaging in HIV-infected and uninfected groups. Data are presented as mean ± SEM, * denotes p<0.05, ** denotes p<0.01, and ns denotes not significant differences.

To assess white matter integrity, we evaluated MAG and MOG expression in the corpus callosum and cerebellar white matter tracts. MAG expression was significantly reduced in the corpus callosum of HIV-infected animals, but not in the cerebellum (Figure 6A, B). On the other hand, MOG expression was decreased in both the corpus callosum and white matter fiber tract of the cerebellum, suggesting demyelination and axonal damage (Figure 6C, D).

**Figure 6.**
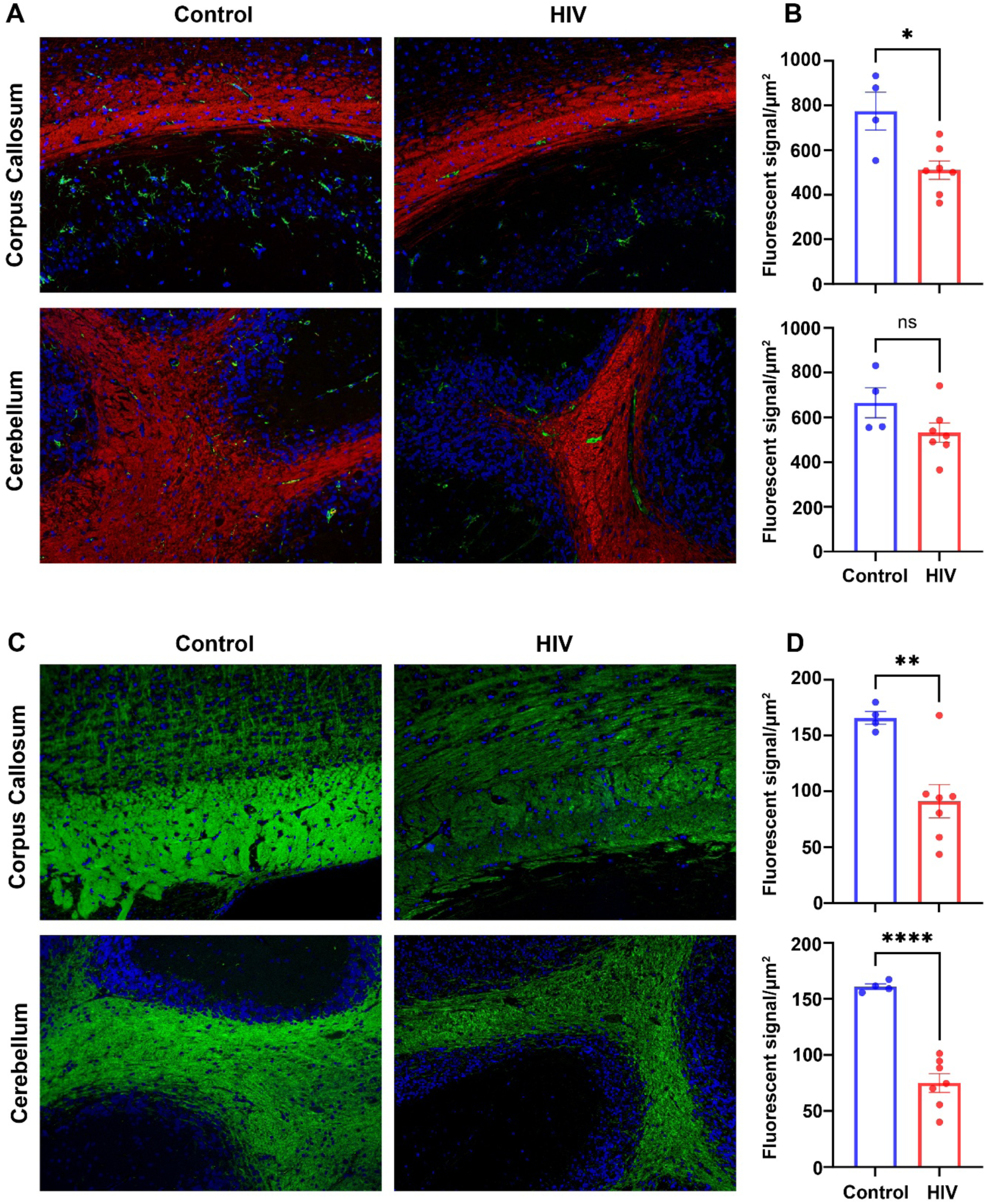
HIV-induced alterations in white matter meylin proteins. Paraffin-embedded 5 µM brain sections were immunostained with myelin antigens **(A)** MAG and **(C)** MOG, along with HLA-DR (green) and nuclear counterstain (DAPI). Representative images acquired at 200x original magnification from the corpus callosum and cerebellar white matter fiber tracts are shown for uninfected and HIV-infected mice. Quantification of fluorescence intensity for **(B)** MAG and **(D)** MOG across brain regions was performed using multispectral imaging, comparing HIV-infected and uninfected groups. Data are presented as mean ± SEM; *p < 0.05, **p < 0.01, ****p < 0.0001, and ns indicates non-significant differences.

### 3.4. Progressive HIV infection induces human microglial activation and anti-viral responses

To examine region-specific molecular alterations associated with HIV infection, we next performed bulk RNA sequencing on the frontal cortex and hippocampus, two brain regions critically involved in cognitive and behavioral abnormalities associated with (Fig. S1) (Akay et al., 2013; Bogdanova et al., 2010; Israel et al., 2019). Given the chimeric nature of this model, sequencing reads were independently mapped to human and mouse reference genomes to characterize species-specific transcriptional responses. Samples were selected based on HIV viral load and RNA quality metrics (RIN/RQN) (Table S2).

Differential expression analysis of human-aligned transcripts revealed a robust transcriptional response to HIV infection in both brain regions. A total of 1565 genes in the frontal cortex and 1975 genes in the hippocampus were significantly differentially expressed between uninfected controls and HIV-infected animals (corrected significant difference value < 0.05; the complete dataset containing fold change and p-values on Zenodo (https://doi.org/10.5281/zenodo.20723232). In the frontal cortex, 1,391 DEGs (88.9%) were upregulated (Figure 7A, B), while 1,844 DEGs (93.4%) were upregulated in the hippocampus of HIV-infected mice relative to uninfected control mice (Figure 7C, D). However, there was no significant difference in the HIV viral load between the frontal cortex and hippocampus (Fig. S2A). Notably, 809 DEGs were shared between the two regions, indicating a largely conserved human transcriptional response to infection (Fig. S3A).

**Figure 7.**
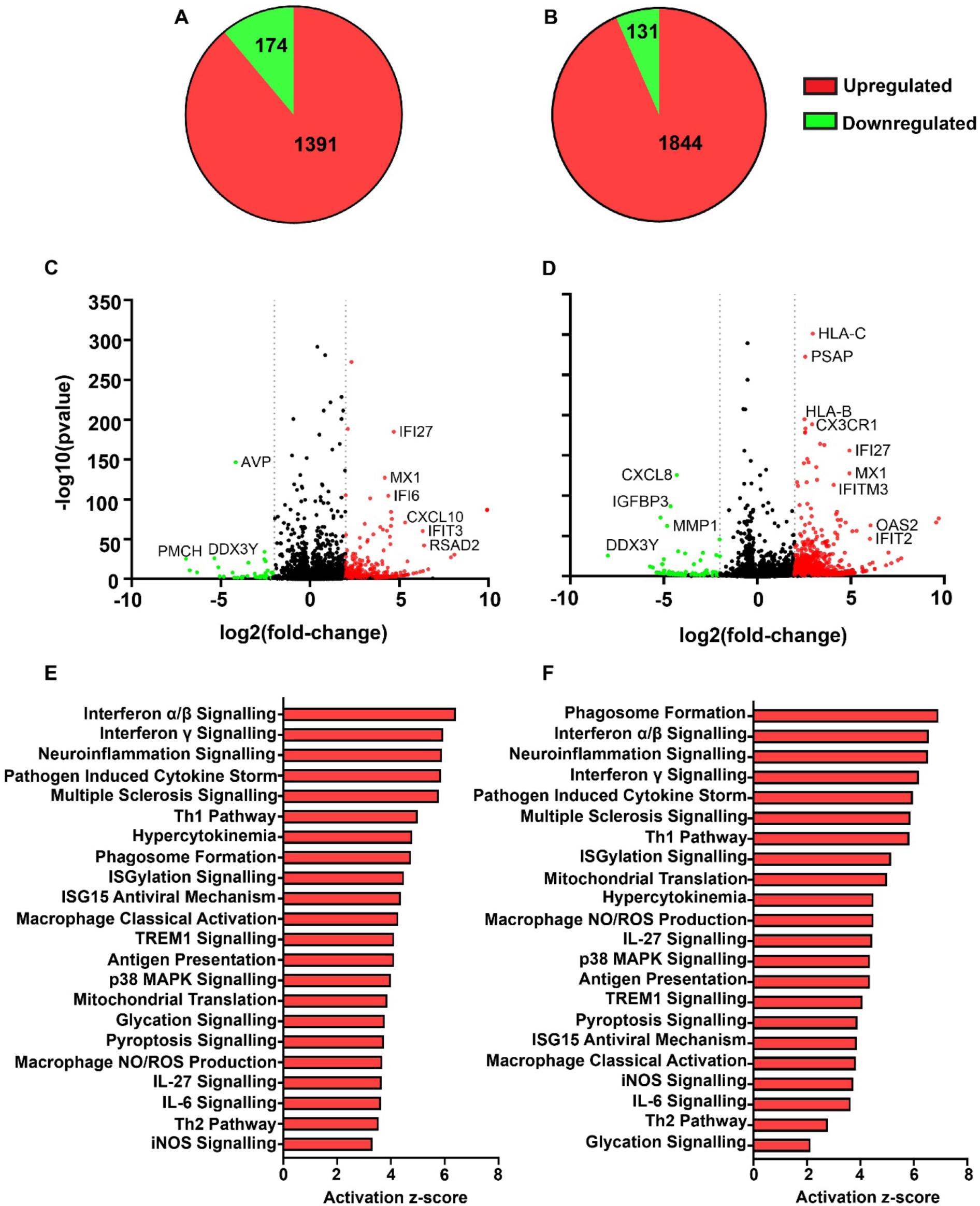
Transcriptional changes in human genes due to HIV-infection. Pie charts of DEGs after alignment of RNA-sequencing reads to the human genome (GRCh37.p13) in comparisons between uninfected and HIV-infected mice in the **(A)** frontal cortex and **(B)** hippocampus. Volcano plots depicting the distribution of DEGs in the **(C)** frontal cortex and **(D)** hippocampus, with significantly upregulated and downregulated DEGs indicated in red and green, respectively. Ingenuity Pathway Analysis of upregulated human genes revealed enrichment of inflammatory and immune-related pathways, including interferon signaling, classical macrophage activation, neuroinflammation signaling, and antigen presentation pathways in the **(E)** frontal cortex and **(F)** hippocampus. Differential expression significance was determined using p-value < 0.05 and with fold-change cutoff of 2.

Pathway enrichment analysis demonstrated that the most significantly upregulated human genes were associated with inflammatory and antiviral processes, including interferon signaling, neuroinflammation signaling, pathogen-induced cytokine storm signaling, and class I MHC antigen processing and presentation pathways (Fig. 7E, F). The upregulated pathways were common to the frontal cortex and the hippocampus. We next performed network analysis using Cytoscape, which revealed extensive interactions among these pathways, identifying highly interconnected hub genes within the DEG network (Fig. 8). Given the substantial overlap in upregulated human immune activation pathways between frontal cortex and hippocampus, the network interaction analysis for the frontal cortex is presented in Figure 8.

**Figure 8.**
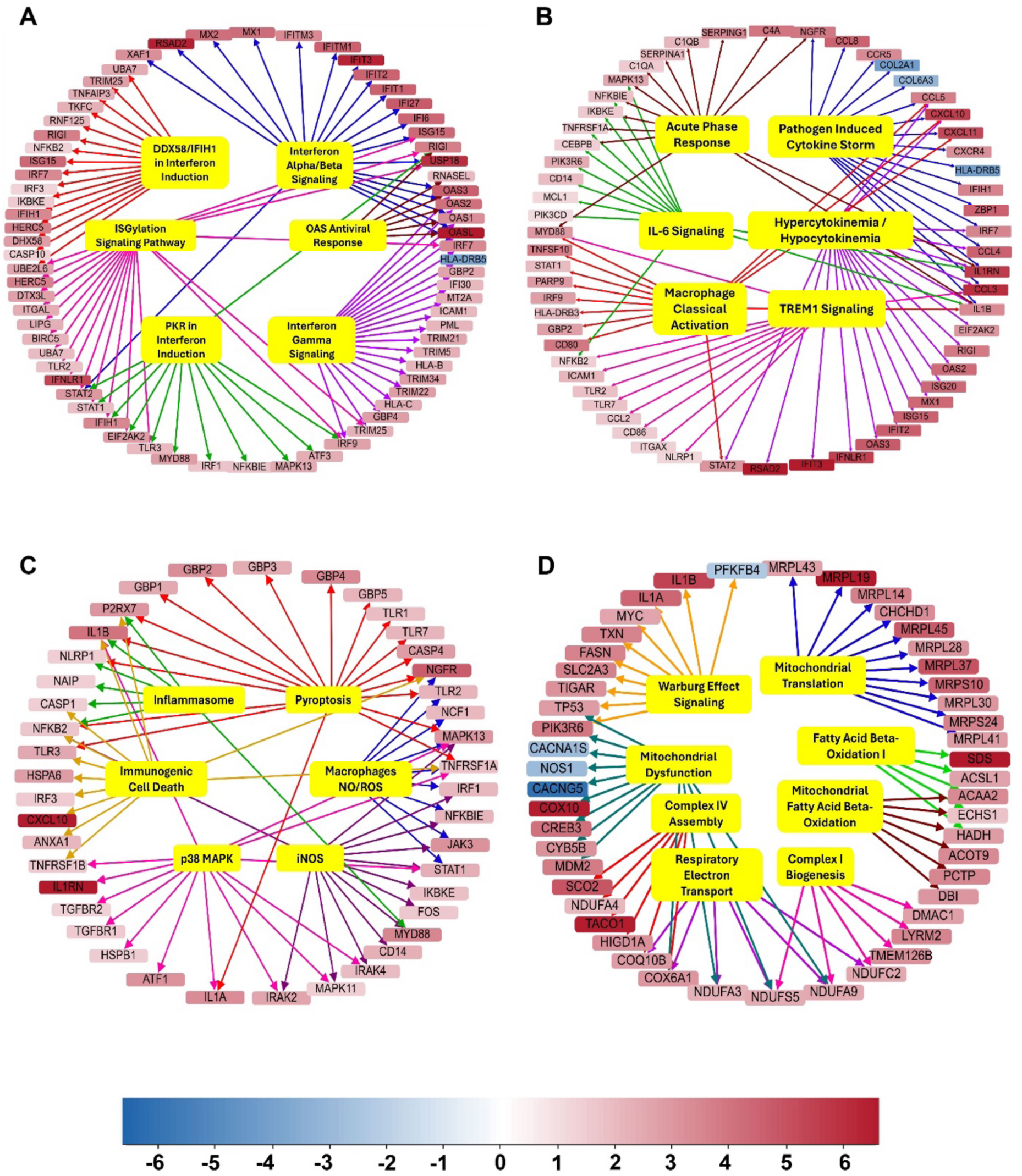
Gene-pathway interaction networks of differentially regulated human genes. Networks were generated using Cytoscape to visualize gene-pathway interactions. **(A)** Antiviral interferon axis (type I/II interferon – ISG antiviral network), **(B)** Neuroimmune activation and cytokine amplification, **(C)** Oxidative stress and inflammatory cell death, and **(D)** Mitochondrial dysfunction and metabolic reprogramming (mitochondrial bioenergetic reprogramming and dysfunction). Nodes represent genes and pathways, and edges indicate their interactions. Upregulated and downregulated genes are highlighted in red and blue, respectively, based on fold change as indicated in the key.

Consistent with predominance in microglia-like human cells in the humanized mouse brain, canonical pathway analysis demonstrated a strong antiviral interferon axis with upregulated type I and II interferon and interferon-stimulated gene (ISG) antiviral network, accompanied by pronounced neuroimmune activation and cytokine amplification. These inflammatory pathways converged on oxidative stress and inflammasome signaling and were associated with mitochondrial bioenergetic remodeling and predicted mitochondrial dysfunction, indicating that progressive HIV infection induces coordinated antiviral and proinflammatory microglial response in both the frontal cortex and hippocampus (Fig. 8A, B, C, D) (Mathews et al., 2019).

### 3.5. Region-specific alterations in murine gene expression reveal neuroinflammatory and neuropathological responses to HIV infection

Given the chimeric nature of the humanized mouse model, we next examined transcriptomic alterations in resident murine brain cells in response to HIV infection and the heightened antiviral and inflammatory responses elicited in human microglia-like cells. Differential expression analysis revealed distinct region-specific responses to HIV infection. A total of 1019 genes in the frontal cortex and 853 genes in the hippocampus were significantly differentially expressed between uninfected controls and HIV-infected animals (corrected significant difference value < 0.05; the complete dataset containing fold change and p-values on Zenodo (https://doi.org/10.5281/zenodo.20723232). In the frontal cortex, most differentially expressed mouse genes were downregulated, with 631 genes (62%) showing decreased expression, whereas the hippocampus exhibited a predominantly upregulated profile, with 730 genes (85.6%) being increased relative to uninfected controls (Fig. 9A, B, C, D). Furthermore, the transcriptional responses were largely region-specific, with only 295 shared genes between the frontal cortex and hippocampus (Fig. S3B).

**Figure 9.**
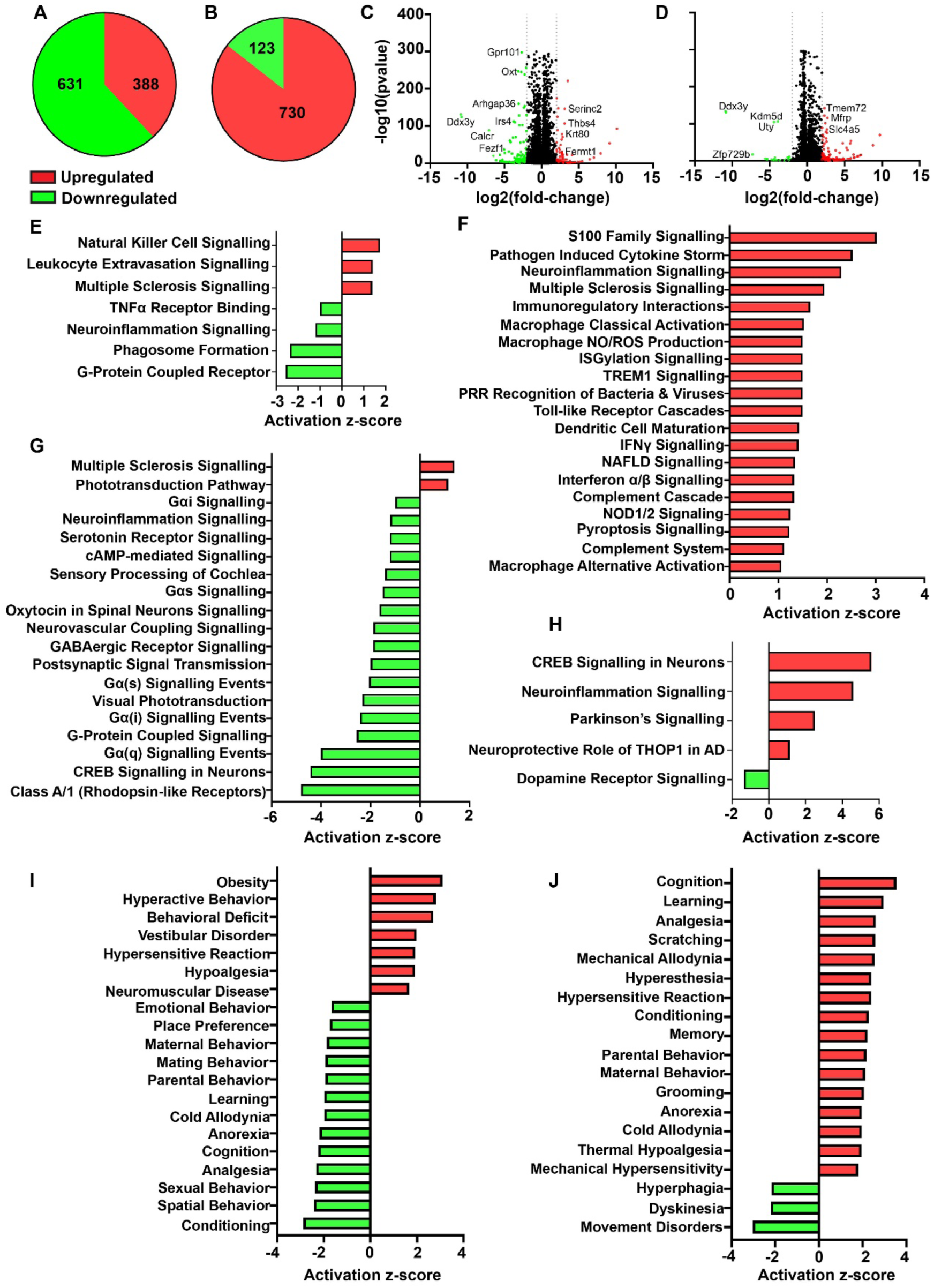
HIV-infection induces transcriptional alterations in mouse genes. Pie chart of DEGs after alignment of RNA-sequencing reads to the mouse genome (GRCm38.p6) in comparisons between uninfected and HIV-infected mice in **(A)** frontal cortex and **(B)** hippocampus. Volcano plot showing gene distribution of DEGs in the **(C)** frontal cortex and **(D)** hippocampus. Ingenuity Pathway Analysis of mouse genes revealed enrichment of mouse inflammatory and immune-related pathways in (**E)** frontal cortex and **(F)** hippocampus. Pathways associated with neurological function are shown in the **(G)** frontal cortex and **(H)** hippocampus. Disease and function analysis identified dysregulated behavioral and neurological functions in **(I)** frontal cortex and **(J)** hippocampus. Upregulated and downregulated DEGs and pathways are represented in red and green, respectively. Differentially expressed genes were identified using a significance threshold of *p* < 0.05 and an absolute fold-change cutoff of ≥ 2.

Pathway analysis showed marked differences in inflammatory and neurological processes between the two regions. Pro-inflammatory pathways were more strongly enriched in the hippocampus compared to the frontal cortex (Fig. 9E, F). In contrast, pathways associated with neurological function and behavior were more prominently dysregulated in the frontal cortex (Fig. 9G, H). Consistent with these findings, transcriptomic analysis predicted greater impairment in cognitive and behavioral functions associated with conditioning, cognition, learning, place preference, and spatial memory in the frontal cortex than in the hippocampus after nine weeks of HIV infection (Fig. 9I, J). To identify genes underlying these behavioral phenotypes, interaction networks between differentially expressed genes and functional pathways were generated using Cytoscape (Fig. 10).

**Figure 10.**
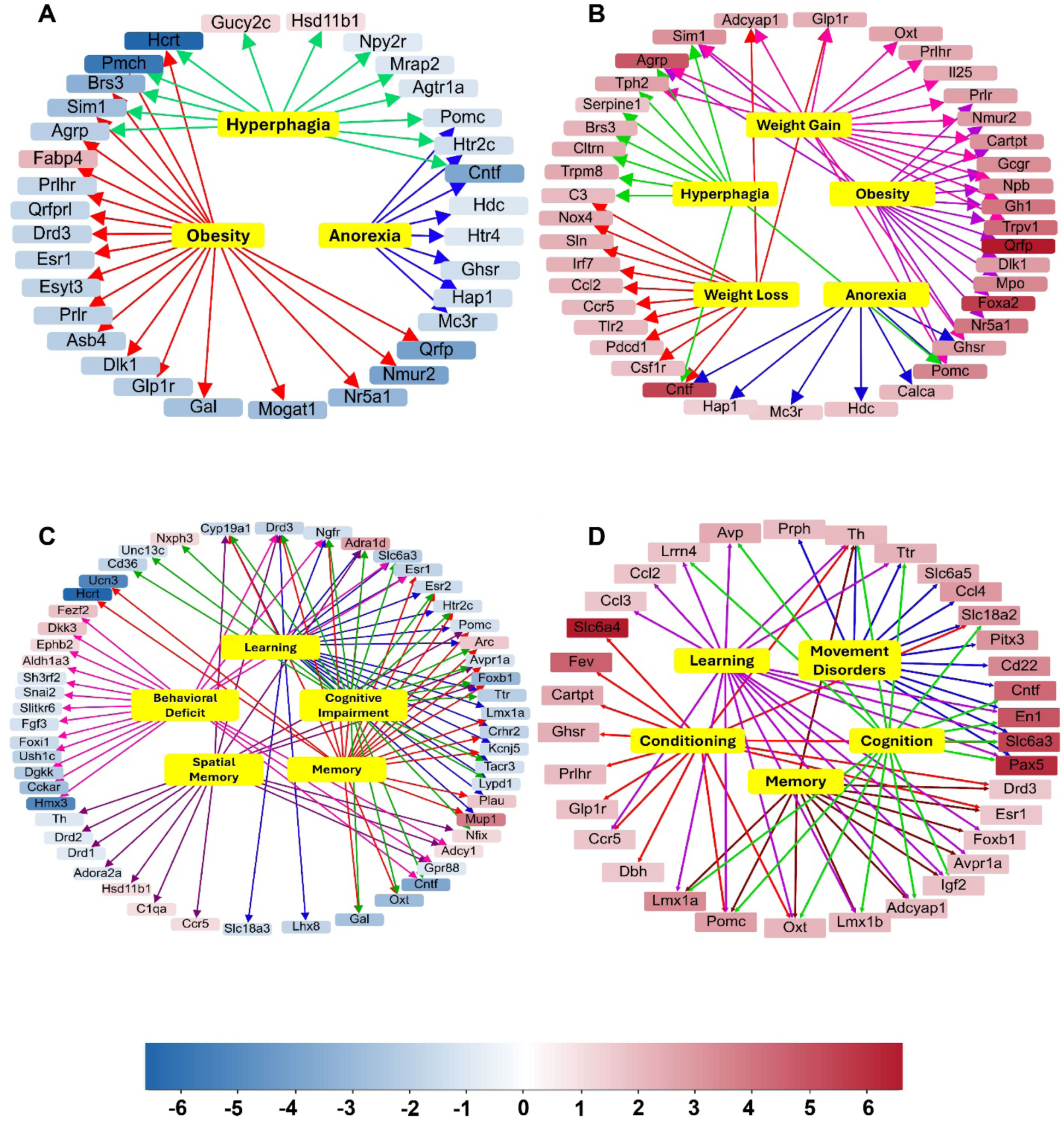
Gene-pathway interaction networks of differentially expressed mouse genes. Networks were generated using Cytoscape to visualize gene-pathway interactions. Feeding-related pathways and genes are shown in the **(A)** frontal cortex and **(B)** hippocampus. HIV-associated gene-pathway Pathways and genes associated with learning, cognition, and behavior are shown in the **(C)** frontal cortex and **(D)** hippocampus. Nodes represent genes and pathways, and edges indicate their interactions. Upregulated and downregulated genes are highlighted in red and blue, respectively, based on fold change, as indicated in the key.

In the frontal cortex, forty-three genes were shared among pathways associated with HAND pathways such as cognition, learning, conditioning, behavioral deficit, cognitive impairment, recognition memory, and spatial memory. These genes included regulators of dopamine neurotransmission (Drd1, Drd2, Drd3), synaptic plasticity (Arc, Egr1), and neuroinflammatory regulators (Ccr5, C1qa, Cntf) (Fig. 10C) (Atluri et al., 2013; Gelman et al., 2012; Jacobs et al., 2013; Kim et al., 2018; Li et al., 2005; Lin et al., 2009; McGuire et al., 2016; Nitkiewicz et al., 2017; Nolan et al., 2019; Riviere-Cazaux et al., 2022; Villalba et al., 2015). In addition, eight genes, including Pomc, Htr2c, Agrp, Sim1, Cntf, Pmch, and Hcrt were associated with feeding-related phenotypes such as anorexia, obesity, and hyperphagia (Fig. 10A) (Deem et al., 2022; Millington, 2007; Mul et al., 2011; Tan et al., 2020; Xi et al., 2012). Similarly, in the hippocampus, thirty-one genes were shared across pathways associated with HAND, including cognition, learning, conditioning, and memory. Many of these genes were specific to neuroinflammation (C3, Ccr5, Ccl2, Ccl3) and dopaminergic signaling (Th and Drd3), suggesting that coordinated alterations in neuroinflammatory and neurotransmitter pathways contribute to HIV-associated cognitive dysfunction (Fig. 10D) (Kuijpers et al., 2010; Lehmann et al., 2019; Liao et al., 2020; Tropea et al., 2024; Tsetsenis et al., 2022). Further, nineteen DEGs linked to food-related behaviors were identified, including key regulators of appetite and energy homeostasis such as Pomc, Agrp, Cartpt, Ghsr, and Glp1r (Hsu, Hahn, Konanur, Lam, et al., 2015; Hsu, Hahn, Konanur, Noble, et al., 2015). These genes were associated with phenotypes including hyperphagia, obesity, anorexia, and body weight regulation, indicating that HIV infection of the CNS disrupts hippocampal pathways involved in metabolic regulation and energy balance (Fig. 10B).

Collectively, these findings demonstrate that HIV infection elicits distinct region-specific transcriptional responses in murine cells and suggest extensive inter-species crosstalk between human microglia and the surrounding mouse neural environment, resulting in differential neuroinflammatory, neurobehavioral, and metabolic consequences in the frontal cortex and hippocampus.

## 4. Discussion

As a substantial number of PLWH on ART experience cognitive impairment and mood disorders despite effective ART, understanding mechanisms underlying HAND remains critical for therapeutic development. The novel humanized glial mouse model in this study provides unique platform for investigating progressive systemic HIV infection, and interaction between the peripheral immune system and the CNS infection. Engraftment of human HSC into NOG mice expressing human IL-34 transgene enables reconstitution of both a functional immune system and human microglia, thereby supporting productive HIV infection, viral dissemination, and establishment of infection within CNS (Honeycutt et al., 2024; Mathews et al., 2019; Zhang et al., 2024). While behavioral phenotype due to HIV was studied using other mouse models, such as transgenic models expressing viral proteins, EcoHIV infection systems, or humanized mice lacking human microglia, these systems do not fully recapitulate productive HIV infection within the CNS (Boska et al., 2014; Kim et al., 2023; Li et al., 2017; Zhang et al., 2023). Nonhuman primate models offer important advantages for studying HIV neuropathogenesis and microglial biology in a physiologically relevant setting, however, their widespread sue is constrained by high costs. limited scalability, and reduced genetic tractability (Hu, 2005; Terrade et al., 2021). To address these limitations, we employed humanized glial mouse model that supports productive HIV infection and allows investigation of behavioral phenotypes and associated neuropathology to model human disease (Mathews et al., 2019). The major strength of this present study is the integration of humanized glial model of HIV brain infection with a multimodal experimental framework that combines long-term home cage behavioral monitoring, detailed neuropathological assessment, and transcriptomic profiling to enable a systems-level understanding of neuroHIV. To our knowledge, this is the first study to utilize extended-duration, noninvasive home cage monitoring in an HIV rodent model to capture spontaneous feeding, drinking, and circadian activity without human intervention. This approach minimizes experimenter-induced stress and provides a more physiologically representative readout of behavioral organization over time. The baseline behavior of uninfected humanized mice was consistent with expected nocturnal activity patterns, including increased feeding and drinking during the dark phase, aligning with established murine circadian psychology (Kobayashi et al., 2020). This supports the validity of the model and provides an important behavioral reference for interpreting infection-associated changes. Such baseline characterization is particularly important in humanized mouse models, where immune reconstitution and xenografting procedures could potentially influence behavioral outcomes independent of HIV infection.

HIV infection was associated with alterations in efficiency and circadian structure, including prolonged feeder engagement without proportional increases in intake and a blunted transition in feeding and drinking behavior at the dark/light cycle boundary. These changes suggest a disruption in the coordination of motivational and circadian regulatory systems. In addition, associations between brain viral burden and increased locomotor activity during the light phase are consistent with virus-linked disruption of rest-activity rhythms and suggest potential sleep/wake dysregulation. These behavioral alterations are consistent with clinical observations in PLWH, in which viral proteins such as Tat have been implicated in circadian disruption and autonomic rhythm dysregulation by decreasing the circadian amplitudes of blood pressure and heart rate (Wang et al., 2014). Similarly, another clinical study reported relationships between viral load and sleep quality measures, further supporting a link between systemic or CNS viral burden and behavioral rhythm integrity (Balthazar et al., 2021).

At the tissue level, neuroinflammatory changes were evaluated across brain regions implicated in executive function, memory, and motor control, including the medial prefrontal cortex, and hippocampus (CA1 and DG). Astrocytic activation was used as the indicator of CNS inflammatory status, given the well-established interactions between astrocytes and microglia in HIV-associated neuroinflammation and neuropathology (Gesualdi et al., 2025; Ton & Xiong, 2013). Given the presence of both human and mouse microglial populations in this model, microglial activation status was further assessed through transcriptomic analyses, with differential expression profiles stratified to distinguish human- and mouse-derived DEGs. The widespread astrocytic response observed across multiple regions is consistent with a state of chronic neuroinflammation, which is known to contribute to synaptic dysfunction, impaired neuronal signaling, and progressive circuit instability. The impact of HIV infection on neuronal myelin integrity, as we observed in humanized mice, was significant across several brain regions. Studies performed on EcoHIV, Tat transgenic, and gp120 models have revealed that while synaptic injury involves both pre- and post-synaptic compartments, microglia preferentially mediate pruning of the pre-synaptic compartments in the frontal cortex, whereas astrocytes increase pruning of both compartments, particularly for excitatory synapses (Ewens et al., 2024; Li et al., 2023; Liang et al., 2025; Watson et al., 2025). Our previous studies have demonstrated that HIV infection in humanized mice without microglia, where HIV-infected human cells were mainly found in meninges and perivascular spaces and rarely in parenchyma, had a significant decrease in expression of myelin proteins MOG and MAG, particularly in the white matter tracts of the brain, such as the corpus callosum (Boska et al., 2014). Similarly, studies in PLWH have reported reductions in synaptic density and dendritic complexity, which correlate more closely with neurocognitive impairment than neuronal loss itself (Everall et al., 1999; Hammond et al., 2018; Masliah et al., 1997; W. Ru & S.-J. Tang, 2017; Sá et al., 2004). White matter abnormalities and oligodendrocyte dysfunction have also been documented in both clinical and experimental studies (Liu et al., 2016; Schenck et al., 2024; Zou et al., 2015). Consistent with these observations, analysis of neuronal and myelin markers in the present study supports the conclusion that HIV-associated neuropathology is region-specific rather than uniform throughout the brain. Cortical and cerebellar regions appeared particularly susceptible to synaptic and dendritic injury, whereas the dentate gyrus exhibited evidence suggestive of selective postsynaptic vulnerability. Although hippocampal dysfunction and disrupted neurogenesis are well recognized in both human HIV infection and transgenic mouse models, most previous studies have focused on dendritic abnormalities and postsynaptic dysregulation within the dentate gyrus. In contrast, relatively little is known regarding axonal pathology in this region (Irollo et al., 2021; Lee et al., 2011; Li et al., 2022). Measuring the expression of different myelin proteins revealed the impact of HIV infection on myelin loss and oligodendrocyte dysfunction, with the corpus callosum being vulnerable to both. Collectively, these findings support the notion that HIV-associated CNS disease results from complex interactions among neuroinflammation, synaptic dysfunction, and white matter injury, with distinct brain regions exhibiting differential susceptibility to pathological processes.

We further leveraged humanized glial mouse brain tissues to identify both mouse and human-specific transcriptional responses across multiple brain regions to study the relationship between antiviral and inflammatory responses in relation to behavioral and histopathological findings (Gauthier et al., 2025; Mathews et al., 2019; Ray et al., 2025). This multimodal approach strengthens its translational relevance by providing phenotypic anchors (noticeable behavioral and pathological changes) to changes in molecular signatures in the CNS (Gauthier et al., 2025; Wu et al., 2016). Bulk RNASeq analysis of tissues from humanized brains revealed significant human-specific antiviral ISGs, interferon signaling, macrophage activation, neuroinflammation, and cytokine storm signaling responses, which can be linked to regions of the brain showing significant neuroinflammation as well as neuronal and myelin damage (Gauthier et al., 2025; Honeycutt et al., 2024; Ray et al., 2025). By focusing on both the hippocampus and cortex and employing bulk RNASeq, we achieved robust identification of human antiviral and inflammatory signatures associated with neuropathology. The significant shift towards an inflammatory phenotype, with minimal differences in human gene expression between the regions, suggests that the human microglia from the two regions behaved similarly to clinical HIV infection. Pathway analysis showed that antiviral immune response to HIV infection in the CNS was dominated by type I/II interferon signaling (Boreland et al., 2024; Tang et al., 2025; Thaney & Kaul, 2019). This antiviral response leads to the activation of the resident immune cells, in turn enhancing neuroinflammation and cytokine response. Tissue injury is further amplified by oxidative stress and inflammasome activation (Mamik et al., 2017; Walsh et al., 2014). In parallel, pathway analysis predicted progressive mitochondrial dysfunction and bioenergetic reprogramming, indicating that strain in the cellular energy machinery leads to chronic immune activation, which disrupts glial and neuronal metabolic homeostasis (Ivanov et al., 2016; Ma et al., 2026).

Interestingly, upon analyzing mouse genes, we observed activation of several neuroinflammation-associated pathways, indicating significant crosstalk between human and mouse cells, as the latter are not capable of supporting HIV infection. However, mouse gene expression showed regional differences: pathways specifically related to neuronal injury and behavioral deficits were enriched in the cortex, whereas the hippocampus appeared to be the primary site of inflammatory activity (Honeycutt et al., 2015; Honeycutt et al., 2024; Ray et al., 2025). A similar pattern is observed in primate studies, where the data showed that acute SHIV infection induced a particularly strong neuroinflammatory response, including perturbations in mitochondrial and synaptic pathways, T cell-related inflammation, and microglial activation, specifically in the prefrontal cortex and superior temporal cortex, while changes in the hippocampus were present but less pronounced in the early stage of the infection (Hawes et al., 2022). Transcriptome sequencing of the brain in HIV-transgenic rats showed that immune responses and neurotransmission-related pathways, such as dopamine signaling, growth factor signaling, myelin, and translation-regulatory pathways, are altered in ways that align with impairments in learning, memory, and reward-related behaviors. This occurs in a region-specific manner in the prefrontal cortex, hippocampus, and striatum (Li et al., 2013). Nicotine-rescue experiments utilizing the same transgenic rat model further revealed that the frontal cortex and hippocampal circuits differed not only in their transcriptomic vulnerability to HIV proteins, but also in the molecular programs that support functional recovery, with the cholinergic modulation restoring Wnt / β-catenin and ephrin B signaling in the former and CREB signaling and glutathione metabolism in the latter (Cao et al., 2013).

These findings were consistent with previous research conducted on human patients that demonstrated neuronal damage in the frontal cortex as a hallmark of early stage HIV disease which could result in diminished cortical thickness and general grey matter atrophy in turn leading to the disruption of network connectivity in the brain and executive function (e.g., decision making, planning, attention, processing speed, and working memory) (Ipser et al., 2015; Lentz et al., 2009; Ragin et al., 2012; Thompson et al., 2005; Woods et al., 2009). Our findings are corroborated by numerous studies, which have indicated that hippocampal atrophy, a hallmark of cognitive decline and memory deficits, seems to appear much later in the disease course (Maki et al., 2009; Nguchu et al., 2025; Sharma et al., 2022; Thompson et al., 2005). Taken together, this suggests that the frontal cortex exhibits early disruptions in neurotransmission, myelin integrity, and protein synthesis, whereas hippocampal dysfunction appears to be more tightly coupled to late-emerging activity-dependent transcription and alterations in redox balance (Cao et al., 2013; Li et al., 2013). The neuroinflammatory cascade makes the hippocampus particularly susceptible to direct neuronal damage and death (Ferrell & Giunta, 2014; Thompson et al., 2005).

In conclusion, this study has certain limitations that should be considered when interpreting the findings. Variability in human cell engraftment and differences in HIV infection levels across animals may contribute to heterogeneity in the observed outcomes. Future studies with expanded cohorts are expected to help further strengthen the robustness and generalizability of these trends and reduce the impact of such biological variability. Despite these considerations, the data presented here represent an important initial step in validating the NOG-hIL-34 mouse model as a relevant and potentially powerful system for studying HAND in humans. Future investigations could extend these findings by examining sex-dependent behavioral differences, the impact of antiretroviral therapies and drugs of abuse, and by incorporating higher-resolution approaches such as single-cell profiling to resolve distinct cellular populations and states. Overall, this study provides valuable insights into the development and characterization of a functional model of HAND, which can serve as a platform for addressing these critical questions in future.

## Credit authorship contribution statement

**Amanda Fernandes:** Writing original draft & editing, Visualization, Validation, Methodology, Investigation, Formal analysis, Data curation. **Ed Makarov:** Investigation. Methodology, Data curation. **Saumi Matthews:** Investigation, Methodology Data curation. **Debashis Dutta:** Writing-review and editing, investigation**. Matthew Thiele:** Investigation. **Mystera Samuelson:** Investigation. Methodology, Validation, Formal analysis. **Santhi Gorantla:** Writing original draft, review & editing, Visualization, Validation, Supervision, Resources, Project administration, Methodology, Investigation, Funding acquisition, Formal analysis, Data curation, Conceptualization.

## Acknowledgements

We thank UNMC Comparative Medicine Department for the care for animals and their support. We also thank the UNMC Animal Behavior Core (RRID:SCR 018830) for the Home Cage behavior studies. This work was supported by The National Institute of Health through grants NIH/NIDA 1R01DA054535, NIH/NIMH 1R21NS139920, and NIH/NINDS 1R01MH128009.

