## Supplemental figures and Tables for "HIV-1 Infection in Humanized Microglial Mice Disrupts Feeding Behavior and Circadian Rhythms with Cortical Neuroinflammation"

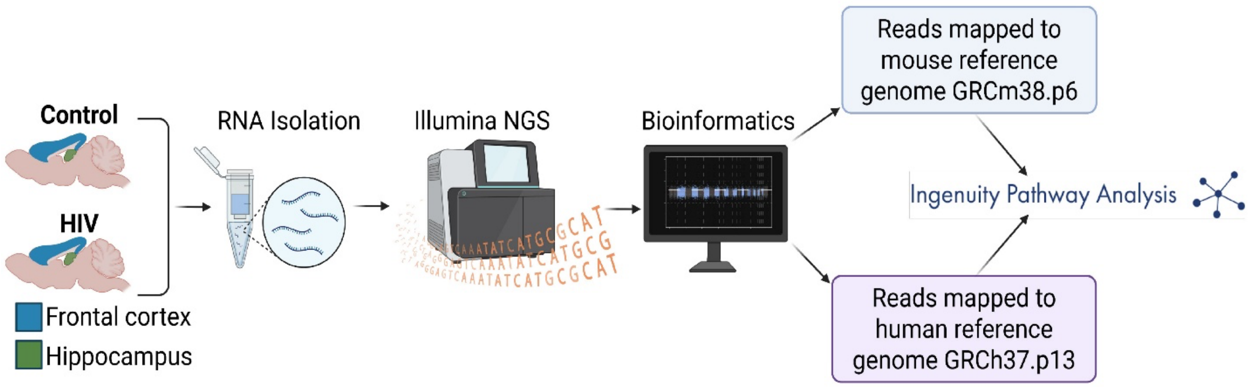


**Supplementary Figure S1.** Experimental schematic illustrating brain RNA isolation and bulk RNA-seq analysis.

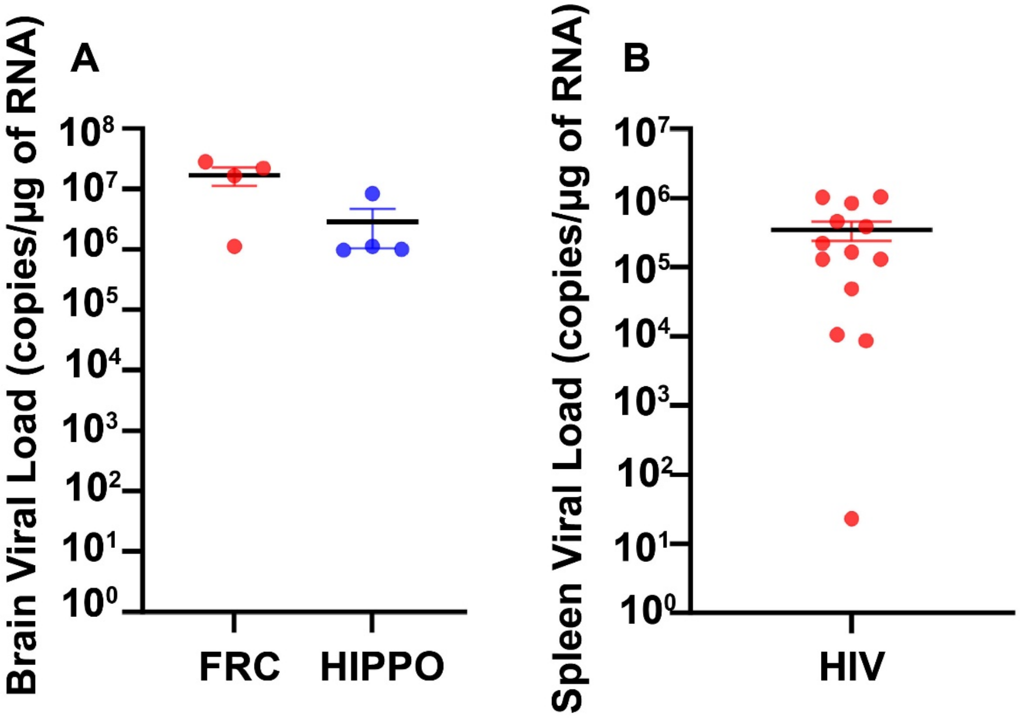


**Supplementary Figure S2. HIV infection levels in brain regions.** HIV viral RNA levels measured by ddPCR **(A)** in the frontal cortex (FRC) and hippocampal (HIPPO) regions of the brain from HIV-infected mice used for RNA-sequencing. **(B)** HIV-1 RNA levels in the spleens of all mice used for home cage monitoring.

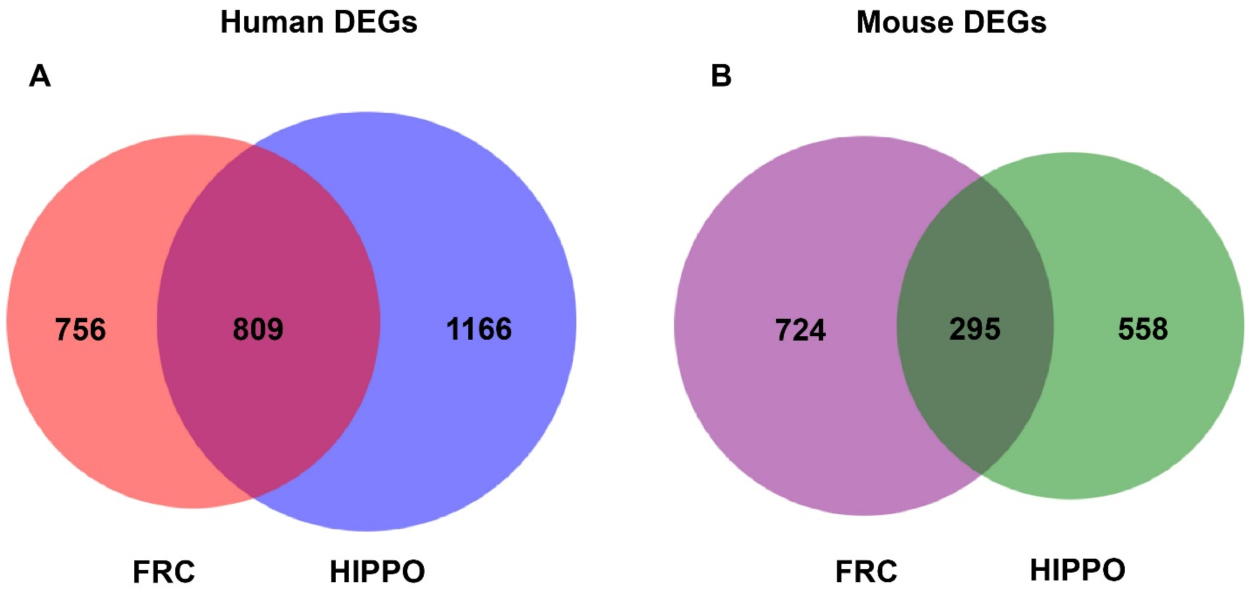


**Supplementary Figure S3.** Venn diagram showing shared and unique **(A)** human genes (red – frontal cortex, blue – hippocampus) and **(B)** mouse genes (purple – frontal cortex, green – hippocampus).

| | | Pre-infection | | | |
|---|---|---|---|---|---|
| **Animal ID** | **Condition** | **CD45+** | **CD4+** | **CD8+** | **CD14+** |
| 5 | Control | 7.25 % | 75.2 % | 22.2 % | 2.29 % |
| A103 | Control | 18.9 % | 54.5 % | 43.2 % | 7.03 % |
| A105 | Control | 39.5 % | 47.8 % | 46.9 % | 3.54 % |
| A106 | Control | 52.6 % | 83.9 % | 14.9 % | 17.7 % |
| A107 | Control | 25.2 % | 52.3 % | 40.9 % | 4.56 % |
| B852 | Control | 38.3 % | 81.8 % | 14.6 % | 3.68 % |
| B855 | Control | 15.6 % | 68.4 % | 28.8 % | 3.99 % |
| B978 | Control | 9.42 % | 72.0 % | 21.4 % | 6.20 % |
| B979 | Control | 13.3 % | 56.5 % | 40.3 % | 0 % |
| B980 | Control | 24.6 % | 63.0 % | 32.3 % | 3.53 % |
| B995 | Control | 14.4 % | 66.5 % | 33.1 % | 0 % |
| 1 | HIV | 21.6 % | 79.0 % | 18.0 % | 1.68 % |
| 3 | HIV | 35.4 % | 80.8 % | 15.9 % | 1.92 % |
| 4 | HIV | 45.1 % | 69.2 % | 27.5 % | 2.45 % |
| A108 | HIV | 18.6 % | 50.0 % | 50.0 % | 6.29 % |
| A111 | HIV | 14.2 % | 68.4 % | 30.7 % | 2.69 % |
| B863 | HIV | 15.3 % | 72.0 % | 23.8 % | 2.90 % |
| B865 | HIV | 43.9 % | 72.7 % | 24.0 % | 4.98 % |
| B872 | HIV | 45.1 % | 69.2 % | 27.3 % | 4.22 % |
| B984 | HIV | 35.2 % | 71.1 % | 23.4 % | 0.63 % |
| B985 | HIV | 20.9 % | 71.8 % | 25.0 % | 0.36 % |
| B989 | HIV | 57.2 % | 70.3 % | 20.6 % | 1.95 % |
| B990 | HIV | 26.1 % | 74.0 % | 21.2 % | 2.93 % |
| B991 | HIV | 39.6 % | 73.9 % | 16.6 % | 6.42 % |
| B998 | HIV | 49.8 % | 75.7 % | 19.4 % | 3.07 % |

**Supplementary Table S1.** Flow cytometry data for mice used in home cage monitoring analysis from peripheral blood before HIV infection.

| Sample Name | Group | Total Mass (µg) | RIN/RQN | 28S/18S |
|---|---|---|---|---|
| 662 Frontal Cortex | Control | 2.88 | 7.1 | 1.2 |
| 662 Hippocampus | Control | 2.912 | 7.4 | 1.3 |
| 671 Frontal Cortex | Control | 1.995 | 5.8 | 0.7 |
| 671 Hippocampus | Control | 2.325 | 6.5 | 1 |
| 688 Frontal Cortex | Control | 2.655 | 7.7 | 1.3 |
| 688 Hippocampus | Control | 1.034 | 8.7 | 1.2 |
| 690 Frontal Cortex | Control | 2.28 | 7.9 | 1.4 |
| 690 Hippocampus | Control | 2.28 | 7.9 | 1.5 |
| 692 Frontal Cortex | HIV | 3.36 | 6.6 | 1.1 |
| 692 Hippocampus | HIV | 2.07 | 6.8 | 1 |
| 681 Frontal Cortex | HIV | 2.445 | 7.5 | 1.4 |
| 681 Hippocampus | HIV | 2.535 | 8.5 | 1.7 |
| 679 Frontal Cortex | HIV | 2.355 | 6.8 | 0.9 |
| 679 Hippocampus | HIV | 1.965 | 6.8 | 1 |
| 674 Frontal Cortex | HIV | 2.895 | 8.3 | 1.6 |
| 674 Hippocampus | HIV | 1.89 | 8.4 | 1.6 |

**Supplementary Table S2.** Quality assessment of RNA samples used for RNA sequencing.
